# Lamina cribrosa vessel and collagen beam networks are distinct

**DOI:** 10.1101/2021.10.03.462932

**Authors:** Susannah Waxman, Bryn L. Brazile, Bin Yang, Alexandra L. Gogola, Yi Hua, Po Lam, Po-Yi Lee, Andrew P. Voorhees, Joseph F. Rizzo, Tatjana C. Jakobs, Ian A. Sigal

## Abstract

Our goal was to analyze the spatial interrelation between vascular and collagen networks in the lamina cribrosa (LC). Specifically, we quantified the percentages of collagen beams with/without vessels and of vessels inside/outside of collagen beams. To do this, the vasculature of six normal monkey eyes was labelled by perfusion post-mortem. After enucleation, coronal cryosections through the LC were imaged using fluorescence and polarized light microscopy to visualize the blood vessels and collagen beams, respectively. The images were registered to form 3D volumes. Beams and vessels were segmented, and their spatial interrelationship was quantified in 3D. We found that 22% of the beams contained a vessel (range 14% to 32%), and 21% of vessels were outside beams (13% to 36%). Stated differently, 78% of beams did not contain a vessel (68% to 86%), and 79% of vessels were inside a beam (64% to 87%). Individual monkeys differed significantly in the fraction of vessels outside beams (p<0.01 by linear mixed effect analysis), but not in the fraction of beams with vessels (p>0.05). There were no significant differences between contralateral eyes in the percent of beams with vessels and of vessels outside beams (p>0.05). Our results show that the vascular and collagenous networks of the LC in monkey are clearly distinct, and the historical notions that each LC beam contains a vessel and all vessels are within beams are inaccurate. We postulate that vessels outside beams may be relatively more vulnerable to mechanical compression by elevated IOP than are vessels shielded inside of beams.

**Research highlights:**

- We combined fluorescence and polarized light microscopy to map in 3D the lamina cribrosa vessels and collagen beams of three pairs of monkey eyes
- Collagen beam and vessel networks of the lamina cribrosa have distinct topologies
- Over half of lamina cribrosa collagen beams did not contain a blood vessel
- One fifth of blood vessels in the lamina cribrosa were outside collagen beams
- Beams with/without vessels and vessels inside/outside beams may respond differently to IOP

## 1. Introduction

Both biomechanics and hemodynamics are central to the physiology of the optic nerve head in health and disease. In the lamina cribrosa (LC) in particular, biomechanical as well as vascular factors have been proposed to contribute to the susceptibility to several potentially blinding neurodegenerative diseases such as glaucoma and non-arteritic anterior ischemic optic neuropathy (NAION).^1–13^ Experimentally, biomechanical and vascular factors are often considered separately and independently. However, the close relationship between the collagenous and vascular networks within the LC suggests that these two factors likely have substantial effects on one another, and therefore they have to be considered simultaneously. Moreover, it is essential to understand the relationship between the collagenous and vascular networks of the LC to fully understand their roles in susceptibility to disease.

Current understanding of the LC blood supply is still largely rooted in several classic studies of ONH vasculature and structure.^5,12,14,15^ Based on these studies, a one-to-one relationship between collagen beams and vessels has been suggested.^16–18^ This is to say that every collagen beam of the LC has been believed to contain a vessel, and similarly, all vessels within the LC have been believed to be enclosed within collagen beams. If this were the case, it would have important implications on the extent to which collagen beams of the LC provide mechanical support to local microvasculature, and with this, the sensitivity of perfusion to changes in intraocular pressure. For instance, the density of vessels would follow the density of collagen beams and the size of LC pores would dictate the maximum distance of neural tissue to its vascular supply. However, if the one-to-one model does not hold, the implications would be different: there may be vessels outside beams, closer to the neural tissues, which may aid in diffusion of oxygen and key nutrients, as well as cell-cell communication necessary for effective neurovascular coupling. Vessels outside beams, however, may not be equally protected mechanically by the collagen and thus may be more sensitive to biomechanical insult, particularly at elevated intraocular pressures (IOP). Collagen beams that contain a vessel may be mechanically different than beams that do not have an opening for a vessel.

To the best of our knowledge, there is very limited direct information about the LC microvascular network in monkeys, and how it relates to the collagen beams or axons that pass through the LC, and the one-to-one model remains unproven. In this study, we aimed to quantify the spatial relationship between the LC vascular and collagen beam networks. Specifically, we quantified the number of collagen beams with vessels, beams without vessels, vessels inside beams, and vessels outside beams. Despite the tremendous advances in imaging of recent years, particularly in optical coherence tomography, simultaneous visualization of the microvasculature and connective tissues of the LC remains out of reach except for small, more superficial regions of the optic disc,^19,20^ which is insufficient for our more comprehensive goals. Hence, we analyzed the LC collagen and vasculature using post-mortem perfusion labeling and a combination of fluorescence and polarized light microscopy (PLM) of cryosections.

## 2. Methods

### Eye procurement and vessel labeling

Six eyes of three healthy adult female rhesus macaque monkeys (*Macaca mulatta*) 12 to 16 years of age were studied. All procedures were approved by the University of Pittsburgh’s Institutional Animal Care and Use Committee (IACUC) and adhered to both the guidelines set forth in the National Institute of Health’s Guide for the Care and Use of Laboratory Animals and the Association of Research in Vision and Ophthalmology (ARVO) statement for the use of animals in ophthalmic and vision research. No live animals were utilized. The monkey head and neck regions were obtained within 30 minutes of sacrifice. The anterior chamber of each eye was cannulated to control IOP throughout the experiment using a saline fluid column. After setting the baseline IOPs (see **Supplementary Figure 1**), we waited 15 minutes to allow for viscoelastic effects to dissipate as done previously^21,22^. A suture was placed on the superior anterior sclera of each eye to serve as fiducial markers for orientation. The carotid arteries on each side of the neck were isolated, their proximal ends ligated, and a polyimide micro-catheter (Doccol Inc., Sharon, MA) threaded into the vessel just beyond the ligature. The vascular bed was washed with PBS until the perfusate was clear. DiI and DiD were used to label vasculature, as done elsewhere.^23–25^ DiI and DiD solutions were freshly prepared from a stock solution of 6 mg/ml DiI or DiD in N,N-Dimethylformamide, diluted to the final concentration of 120 μg/ml in PBS with 5% glucose to adjust osmolarity.^23^ We perfused 100 ml of aqueous DiI solution into each carotid artery at a rate of 5-10 ml per min for 10 minutes. A second PBS wash of 50 ml per carotid artery was perfused through the vessels to remove residual DiI. In 4 of the 6 eyes, IOPs were changed to a second IOP (**Supplementary Figure 1**), followed by another 15 min wait. In the other two eyes the baseline IOP was maintained. In eyes with changed IOP, we then perfused 100 ml of an aqueous solution of DiD into each carotid artery at a rate of 5-10 ml per min for 10 minutes. A third PBS wash of 50 ml PBS was perfused to remove residual DiD. We perfused 50 ml of 10% formalin into each carotid and a second perfusion of 50 ml of 10% formalin after a 15-minute wait. After an additional 15 minutes, the eyes were enucleated, making sure to preserve optic nerves at least 10mm in length from the globe. The IOP control lines were switched from saline to 10% formalin columns with the desired IOP level. To complete the fixation, the eyes were immersion fixed overnight in 10% formalin while IOP was maintained.^26^ Perfusion of eyes with DiI and DiD at different IOPs was done in preparation for future studies regarding the effects of IOP on vessel filling. Because the use of multiple dyes does not alter the results of the current study, and because further work needs to be done before reporting conclusions about the effects of IOP on vessel filling, we do not place focus on IOP effects in this manuscript. For analysis, the segmentations obtained from the different dyes perfused at different pressures were combined into a single vasculature segmentation. More details of this process, including the rationale for using various IOP levels, and the implications on the results, are given in the Image Processing and Analysis section below and in the Discussion.

### Histology and Imaging

The globes were hemisected and the retina examined under a dissecting microscope (Olympus MVX10, Olympus, Tokyo, Japan) using fluorescence to evaluate the perfusions. All eyes had satisfactory perfusions, meaning that there was continuous staining of the vasculature without any discernible dark patches or leaks. The ONH and surrounding sclera were isolated using a 14 mm circular trephine. The tissues were placed in 30% sucrose overnight for cryoprotection, flash-frozen in optimum cutting temperature compound (Tissue Plus, Fisher Healthcare, Houston, TX), and sectioned coronally at 16μm thickness with a cryostat (Leica CM3050S). All data presented in this manuscript comes from sections collected serially, with no missing sections through the LC and without observable regions of damage. Immediately after sectioning, both fluorescence microscopy (FM) and PLM images were acquired of each section using a commercial upright microscope (IX83, Olympus, Tokyo, Japan) to visualize the vessels and collagen, respectively. A 4x strain-free objective (UPLFLN 4XP, Olympus, Tokyo, Japan) was used for both FM and PLM. For PLM we used two linear polarization filters and a quarter wave-plate. For FM, we used filters suited for the excitation/emission profiles of DiI (545/605 nm, Olympus U-3N49004) and DiD (620/700 nm, Olympus U-3N49006). When necessary, mosaicking was used to assemble images of the whole scleral canal. The sections were hydrated and cover-slipped during imaging. Fluorescence signal varied depending on the amount and density of dye in the section. To maximize the visibility of the vasculature, the exposure times were chosen by the microscope operator to maximize visibility of vessels without causing saturation. FM images of each label were acquired separately by using a motorized filter wheel. Imaging parameters were independently optimized for each channel. The collagen fibers are highly birefringent and were fairly easy to distinguish from other tissues using PLM. Although PLM can provide local fiber orientation information, in this work we used only monochromatic retardance images obtained as described previously.^27,28^ It must be noted that our PLM technique does not visualize elastin, an extracellular matrix component of the LC that is important, but less abundant than collagen.^29^ Visualizing elastin would require further efforts, perhaps by IHC^30^ or with multi-photon techniques.^31^ PLM was validated to visualize all collagen and only collagen through comparison with other well-established methods to visualize collagen: picrosirius red staining and second harmonic generation imaging (SHG, **Figure 1, Supplementary Figure 2**). We evaluated the quantitative agreement between measurements obtained using PLM and picrosirius red and SHG. DiI and DiD signal were not obscured by any overlying tissues (**Supplementary Figure 3**). Note that FM and PLM image sets were well co-localized as all images were obtained on the same microscope with the same objective within seconds of each other separated only by a change of filters using the microscope motorized rotation wheel, and after confirming that the filters were properly aligned. Before data collection we verified that images from different modes are aligned. Nevertheless, it should be noted that transformations are stored and handled as 32-bit float variables, and thus their application to rotate and translate discrete images stored as pixels involves approximations. Parfocality of signals was verified in the development of this imaging technique. We confirmed that the dyes were well-centered and focused at the same depth and location as the polarized light signal. Additionally, it must be noted that intravascular perfusion of DiI/DiD labels blood vessels regardless of their size. Future work should consider using a more precise definition, which may potentially require more complex and specific labeling techniques. Image acquisition was done using CellSens (Olympus, Tokyo, Japan).

**Figure 1:**
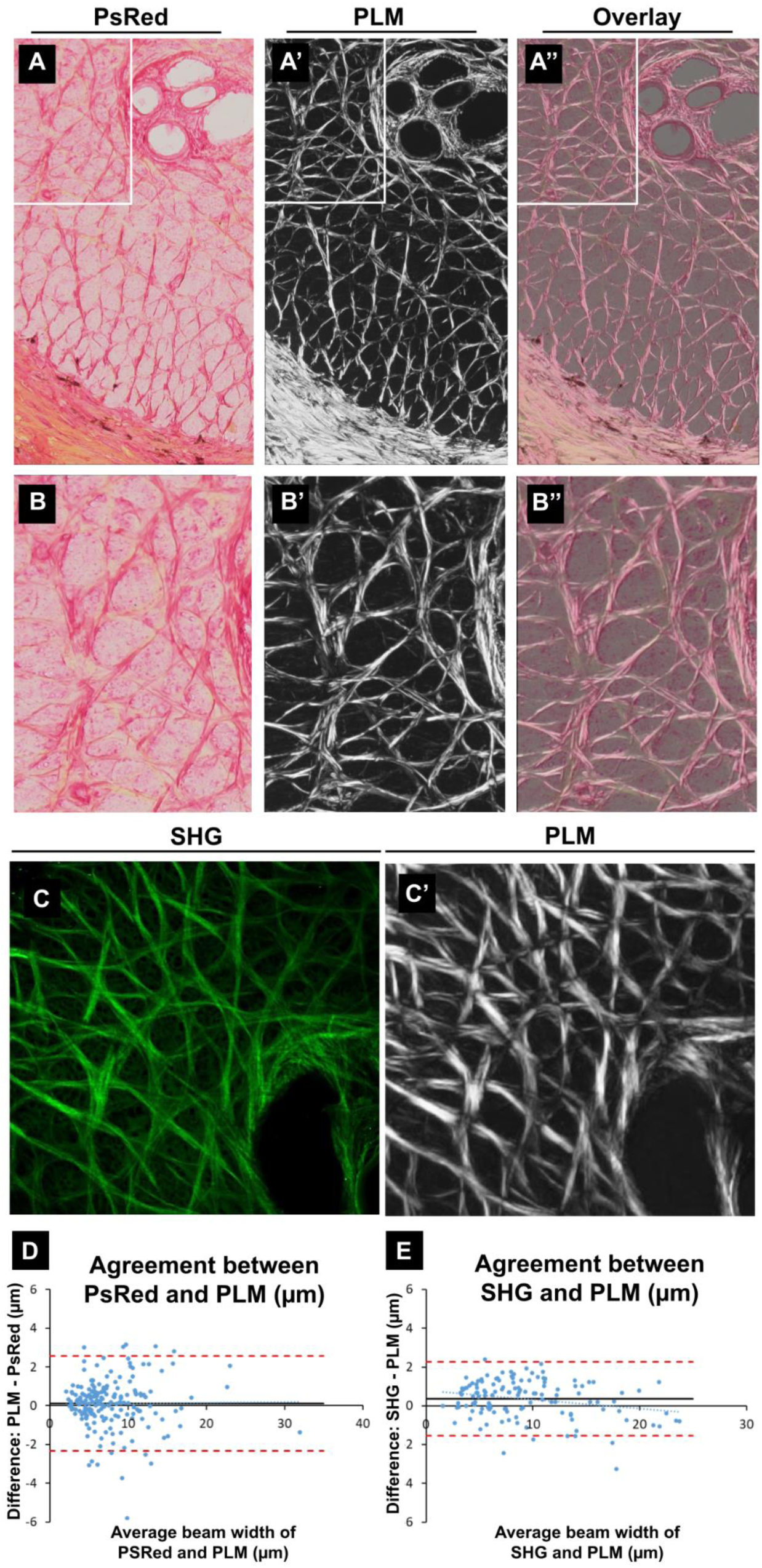
Polarized light microscopy (PLM) detection of LC collagen beams is in line with other well-established methods of collagen visualization. A) A 16µm-thick cryosection through the monkey LC was labeled with picrosirius red (PsRed), a collagen stain, and imaged via brightfield microscopy. Collagen is shown in darker pink/red. A’) The same section imaged via PLM reveals the same pattern of beams visualized via PsRed staining. This is evident in an overlay of these images in A’’. B-B’’ show close up views of the region in A-A’’ within the white box. C) A second harmonic generation (SHG) image of the monkey LC to visualize collagen. C’) The same region of the section in C imaged via PLM as in A’ and B’. The same pattern of beams is visualized among SHG and PLM. The bottom row panels (D and E) show Bland-Altman plots to evaluate the agreement in LC beam widths obtained between PLM and PsRed (left) and between PLM and SHG (right). Overlaid on the scatterpoints on each plot are lines representing the following: best-fit linear function (dotted blue line), mean difference (black line), lower and upper 95% confidence interval limits (dashed red lines). For PLM vs PsRed, the mean difference was 0.12 µm and the SD of the differences was 1.25 µm. For PLM vs SHG, the mean difference was 0.36 µm and the SD of the differences was 0.97 µm. Overall, the plots show excellent agreement between the techniques. Every beam discernible with one technique was also discernible in the other.

### Image Processing and Analysis

Stacks of sequential PLM images were imported, aligned, and registered in Avizo (version 9.1, FEI; ThermoScientific) as previously reported.^32,33^ Briefly, images were registered manually based on landmarks, including tissue edges and marks on the sclera made before sectioning. The translations and rotations were then applied to the FM images, ensuring that PLM/FM image pairs remained perfectly co-localized to each other after stack registration. The FM and PLM images were then merged into red and green channels, respectively, to simultaneously visualize the collagen beam and vessel networks. All analyses were done from reconstructions from stacks of serial 16 µm-thick sections. Projection images were used only to facilitate image visualization and illustration for this manuscript.

In our analyses, LC collagen beams were continuous collagen structures at least 2 pixels wide as detected in our PLM images. Similar definitions have been applied in other investigations of the LC.^34–37^ To mitigate misidentification of continuous collagen beams as non-continuous, analysis was done on 3D reconstructions rather than images of individual sections. This allowed us to detect thin but continuous collagen beams as well as exclude low intensity signal, such as that of collagenous microvascular basement membranes, as well as any imaging noise. It must be noted that cytoskeletal elements of axons also exhibit birefringence, the same property that allows for collagen visualization via PLM. Since cytoskeletal birefringence is orders of magnitude lower than that of collagen fibers, it is generally not difficult to distinguish LC beams from axons, as shown in our images where the pores appear much darker than the beams. Nevertheless, we acknowledge that it could be difficult to distinguish extremely sparse and thin collagen from axons just from the birefringence. In these cases, we can take advantage of the orientation information in PLM.

The vessels and beams were independently segmented from their corresponding PLM and FM images of all sections. LC collagen beams were identified as bright signal under PLM due to collagen birefringence. Networks of linear collagen beams throughout the LC create pores which are filled with retinal ganglion cell axon bundles. Due to the curved shape of the lamina, some sections contained only partial LC (**Fig. 3A**). Hence, the stack of sections is necessary to identify and characterize the full LC. From the beams we then defined the LC region in 3D based on the presence of collagenous beams^38^. Thus, the region we call the LC is equivalent to what is sometimes referred to as the scleral lamina to distinguish it from a more anterior glial or choroidal lamina^39,40^. The analyses of beam and vessel inter-relationship were then done on the LC region, thus excluding the prelamina, retrolamina, and any tissues peripheral to the LC. We first applied a Jerman filter^41^ in Fiji to enhance the contrast of both beams and vessels. We then utilized automated and manual segmentation techniques iteratively in Avizo until a single experienced operator (B.B.) considered all segmentations to be high quality representations of the images. Collagen and each vessel were segmented separately and independently. The vessel segmentations or “labels” were then combined via union to create a single 3D map of the vasculature for each eye. Perfusion with DiI and DiO at different IOPs was done in preparation for future studies in which we intend to assess the effects of IOP on vessel filling. Our goal in this study was directed at a more fundamental understanding of the inter-relationship between vessels and beams. For simplicity, we focused the analysis on these general aspects that are not affected by the use of two dyes.

Following segmentation, we generated distance maps measuring the minimum Euclidean distance between vessel pixels and beam pixels. A distance of zero indicated pixels of overlapping beams and vessels. In these overlapping regions, beams were considered to contain vessels and were therefore counted as regions of *beams with vessels*. Distances of greater than zero indicated regions of *vessels outside beams*. Next, we generated skeleton images of the collagen beams, vessels, and regions of overlap using parallel medial axis thinning.^42^ The process of skeletonization compensates for the variable width of collagen beams. Voxel-by-voxel analysis, rather than beam-by-beam or vessel-by-vessel analysis, compensates for variable lengths of beams and vessels, and for potential multiple crossings. We computed the ratio of *vessels without beams* (R_1_) and ratio of *beams with vessels* (R_2_) for all sections in the central LC as follows:

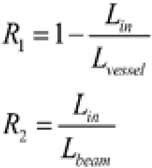

where L_in_, L_vessel,_ and L_beam_ are the skeleton length of overlapping regions, vessels, and beams, respectively. Percentage lengths of *beams without vessels* and *vessels inside beams* were calculated by subtracting the percentage length of *beams with vessels* and *vessels outside beams*, respectively, from 100%. Analysis was done in 3D, as has been proven essential for the complex shape of the LC region.^38^

The use of PLM and FM images to define LC beams and vessels carries some unavoidable uncertainty. To understand how this may affect our results we did a sensitivity analysis. Specifically, we evaluated the effects of systematic changes in beam and/or vessel width on the number of vessels inside/outside beams, and beams with/without vessels (**Figure 2**). As we will show later, the results indicated only a minimal sensitivity of the outcomes, and therefore we only did this study in the OD eye of Monkey 2.

**Figure 2:**
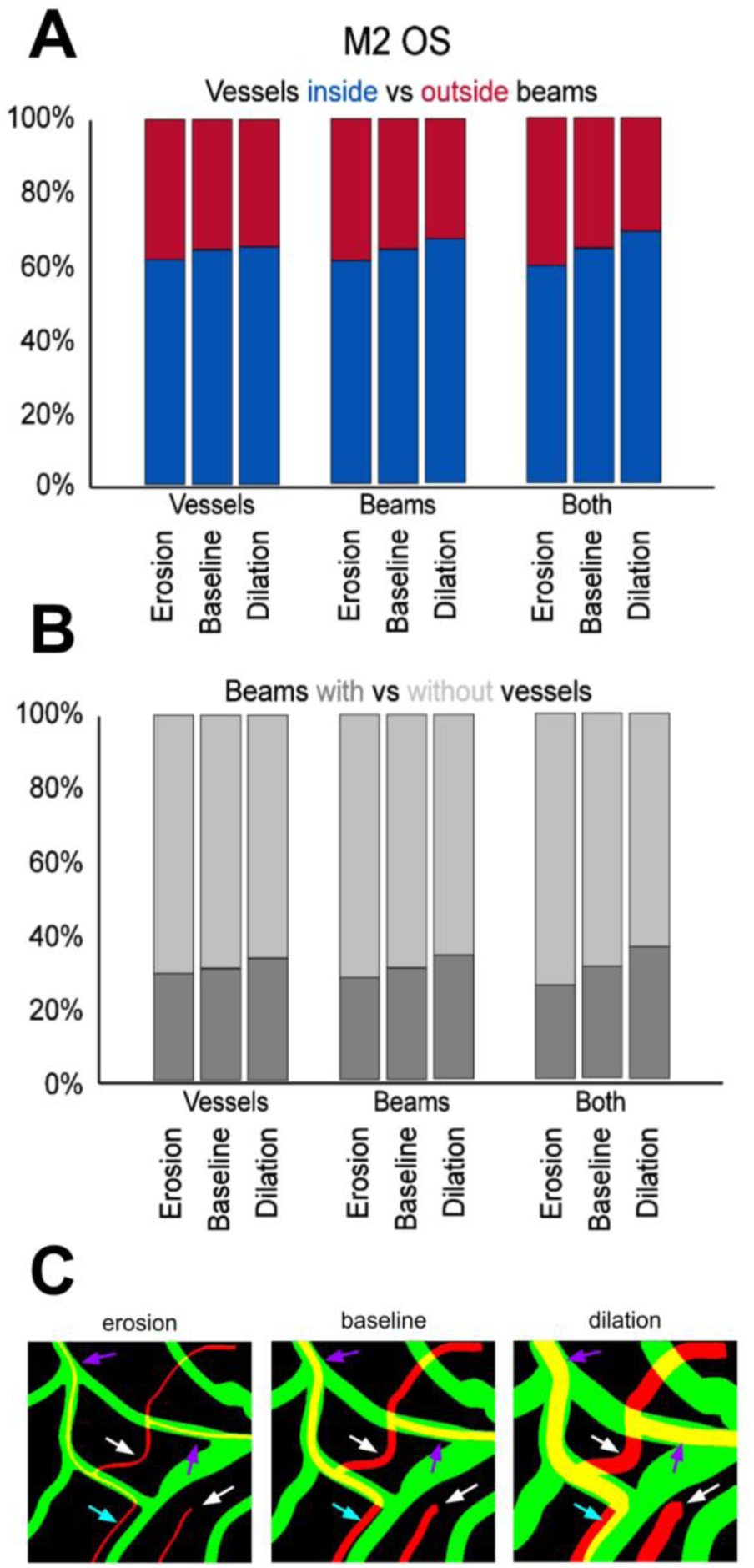
Results from the analysis of sensitivity to LC beam width in the OS eye of M2. The fractions of vessels inside vs outside beams (A) and of beams with vs without vessels (B) were calculated before and after changes (erosion of 3µm or dilation of 6µm) of beams, vessels or both. A visual representation of how erosion/dilation of beams (green) and vessels (red) can affect spatial relationships is shown in the diagram in C. Erosion/dilation of vessels and beams does not alter whether most beams are with/without vessels or whether most beams are inside/outside beams, as shown quantitatively in A and B. White arrows: vessels outside beams that stay outside beams despite erosion/dilation, purple arrows: vessels inside beams that remain inside beams despite erosion/dilation, cyan arrows: vessel outside beam that may appear as inside beam or outside beam depending upon erosion/dilation.

#### Statistics

Correlation between *beams with vessels* and *vessels outside beams, beams with vessels* and individual animal, *beams with vessels* and eye (OD vs. OS), *vessels outside beams* and individual animal, and *vessels outside beams* and eye (OD vs. OS) were assessed through linear mixed effects models fit by maximum likelihood, implemented in R.^43^ Eye (OS. vs. OD), individual animal, and/or histological section were considered to contribute random effects. Findings were considered statistically significant at an alpha < 0.05.

## 3. Results

Comparing LC beam width measurements between PLM and PsRed and PLM and SHG revealed excellent agreement between the techniques (**Figure 1**). Mean LC beam width differences (estimated bias) for PLM vs PsRed was 0.12 µm and the SD of the differences was 1.25 µm. For PLM vs SHG, the mean difference was 0.36 µm and the SD of the differences was 0.97 µm. Overall, the plots show excellent agreement between the techniques. The fit slopes indicate that there were no proportional biases between techniques. A sweep through the Box-Cox family of transformations indicated that no transformation would substantially improve the heteroscedasticity.

The sensitivity analysis showed that there were only minimal changes in the results when the width of the beams, vessels or both were artificially changed (**Figure 2**). The ranges over which we did the sensitivity test (erosions of 3µm and dilations of 6µm) are fairly large compared with the variability observed in the agreement evaluations (**Figure 1D & E**). This is evidence that our choice of imaging or analysis method is not the determining factor of our findings.

PLM allowed us to capture information about the location, density, and orientation of collagen beams of the LC in 16µm-thick sections (**Fig. 3A**). DiD/DiI perfusion and FM allowed us to map vasculature within the same sections (**Fig. 3B**). Individual PLM images were successfully registered and aligned to their serial neighbors to render image stacks of the complete LC. As FM and PLM images were taken within seconds of each other and movement of the sample was not required between this switch in imaging modalities, well-aligned stacks of FM images were successfully created by applying transformations used to align PLM images. Comprehensive maps of LC collagen (**Fig. 3C**) and vasculature (**Fig. 3D**) were readily visualized as depth-resolved maximum intensity projections. We obtained well-registered image stacks of collagen and microvasculature in the ONH, including the LC, for all six eyes.

**Figure 3.**
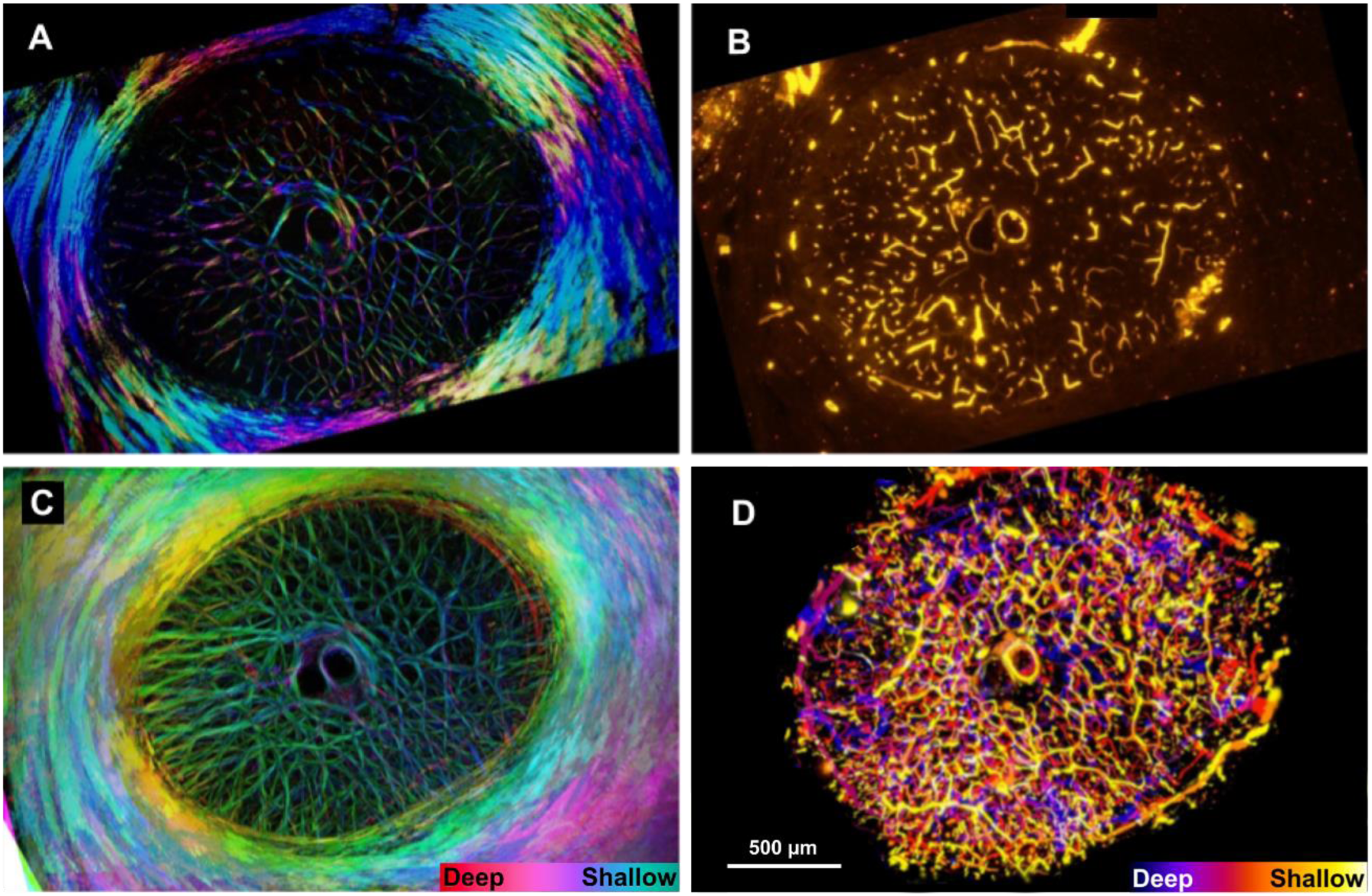
Collagen beams and vessels have distinct topologies across the LC. Example images of collagen (A) and vessels (B) in a single 16 µm-thick coronal section through the ONH at the level of the LC. In A, the color of the collagen is derived from its local orientation, and local intensity of the image is derived from the collagen density. Images of collagen were acquired using PLM (A, C), and images of vessels were acquired using FM (B, D.) C and D show maximum intensity projections of stacks of 19 sections (in C) and 8 sections (in D). Fewer sections were projected in D to avoid overcrowding. In C and D, collagen and vessels are color-coded by depth (see scale at bottom right for each image). Our imaging and segmentation focused on the vessels within the canal. This could cause vessels outside to appear discontinuous or “broken”. Hence, in D, vessels outside the scleral canal have been digitally masked. Contrast was adjusted to best display imaged features.

In individual 16µm sections, both collagen beams and vessels were observed throughout the depth of the sample (**Figure 4**). Upon observation of these merged images, we noticed that the vessels and beams had different topologies across the lamina. We saw that not all vessels were contained within the center of the lamina beams. Instead, we observed: beams that had no vessel at all; vessels that were along the side of and even crossing through adjacent beams; vessels that crossed the lamina pores without any beam support; and consistent with historical belief, vessels residing within collagen beams (**Fig. 5**). We verified these findings in individual 16µm sections (**Fig. 4, 5**), stacks of two sequential images (32 µm-thick, **Fig. 6**), and in whole LC projections (**Fig. 7**). This allowed us to easily discern details as well as a gestalt of vessels and collagen. Even when viewing a maximum intensity projection of the complete LC, these key features were retained. As is usual, the fluorescent intensity of the dyes exhibited some regional variations. While vasculature in some regions was less bright than others, no significant portions of dye-free LC were observed, indicating complete or near-complete vascular perfusion. The analyses were done based on the presence of vessels, and not on their fluorescence intensity. Segmentation of collagen beams and vessels allowed for 3D reconstruction of collagen beams and vasculature (**Fig 8-12**). The cupped shape of the LC region (**Fig. 8**) as well as the interconnected nature of collagen beams and vessels was preserved in these 3D reconstructions (**Fig. 8-12**).

**Figure 4.**
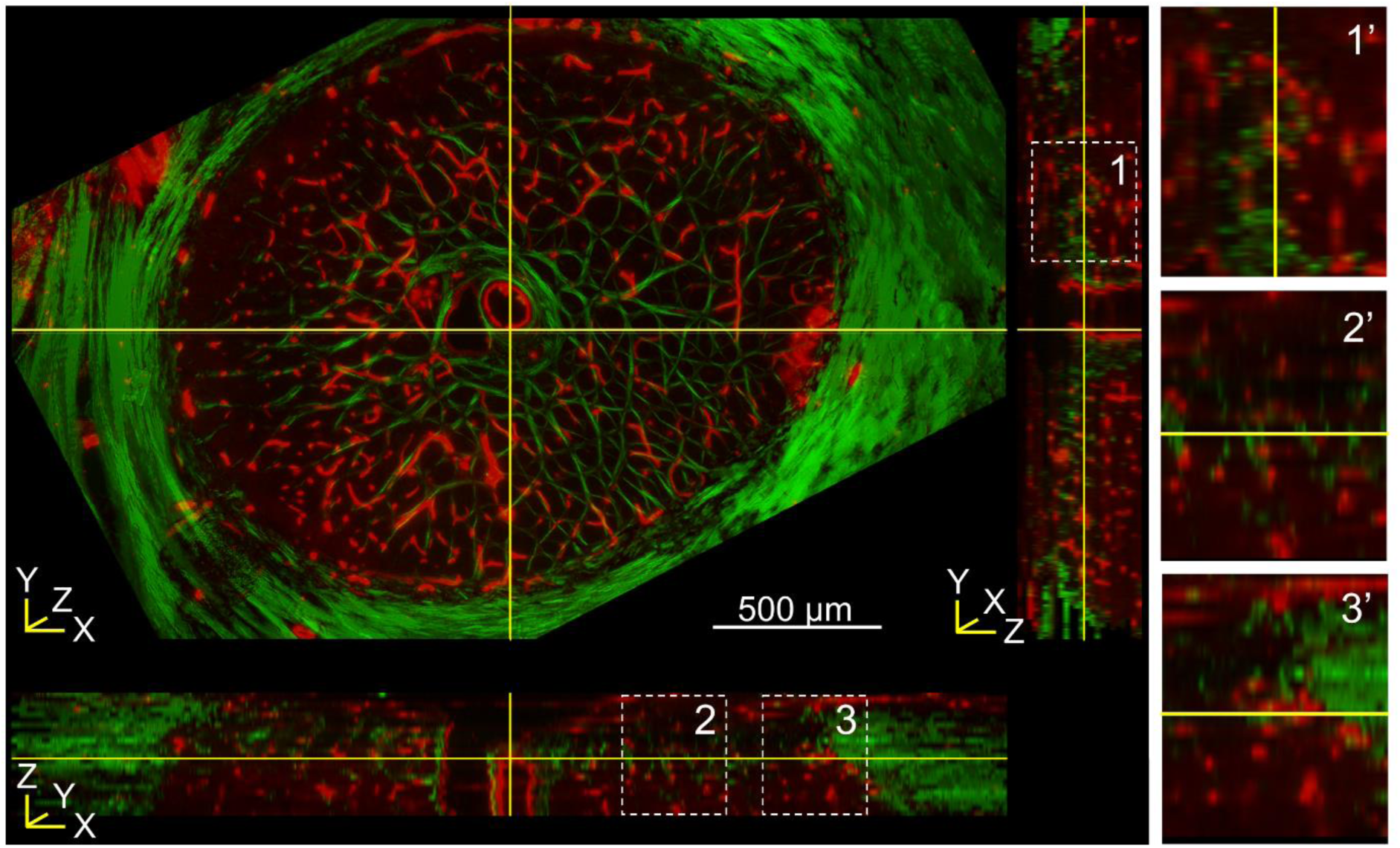
Aligned, registered, and merged FM and PLM images. Red (vessels) and green (collagen) merged image of a single 16 µm -thick coronal section shown along the XY plane. Here, the central retinal artery is well highlighted in the center of the image. Virtual sections along the XZ (bottom) and YZ (right) planes are depicted by the yellow demarcations. Dashed boxes indicate insets which correspond with magnified panels (right) showing detail. In this representative sample, collagen and vasculature were visible throughout. Note that the LC demonstrates a cupped shape, discernible in the coronal section, with beams present in the center and not in the upper left of this example, and in the virtual cross-sections.

**Figure 5.**
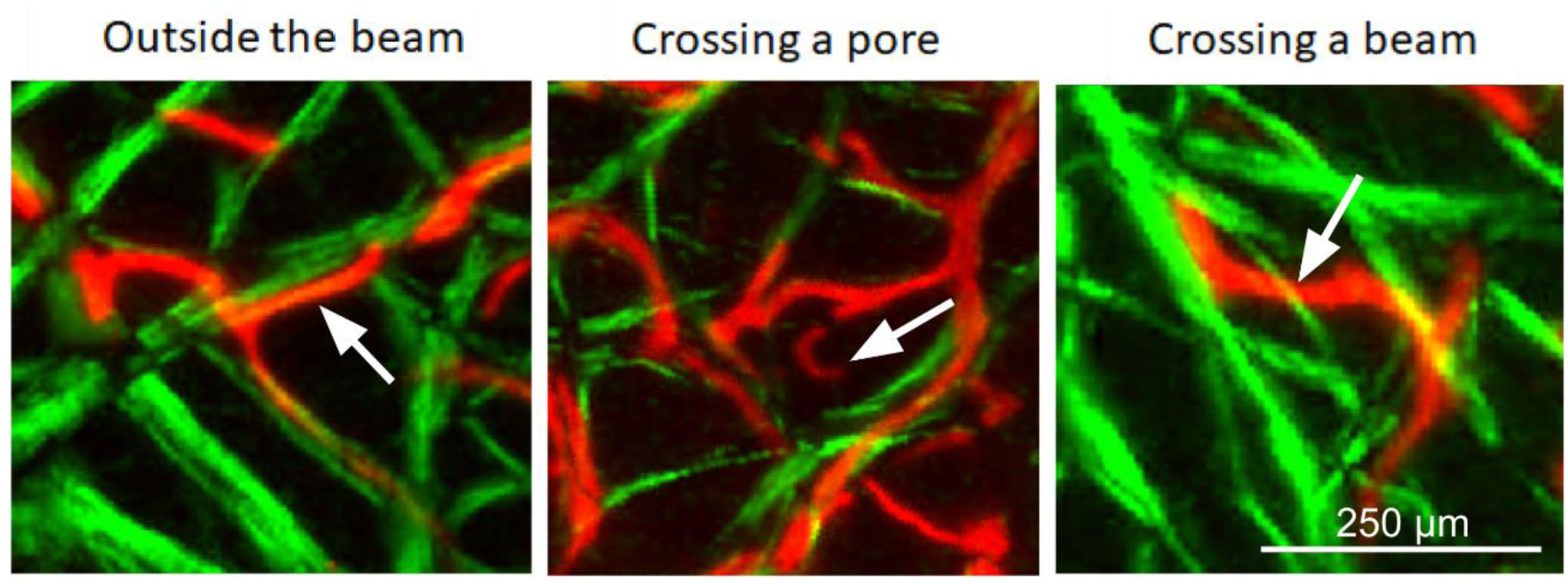
Red (vessels) and green (collagen) merged images of a single 16µm-thick section. Left) Example of a vessel running along the outside of the collagen beam. Middle) Example of a vessel crossing a pore without any collagen support. Right) Example of a vessel that crosses perpendicular to a collagen beam.

**Figure 6.**
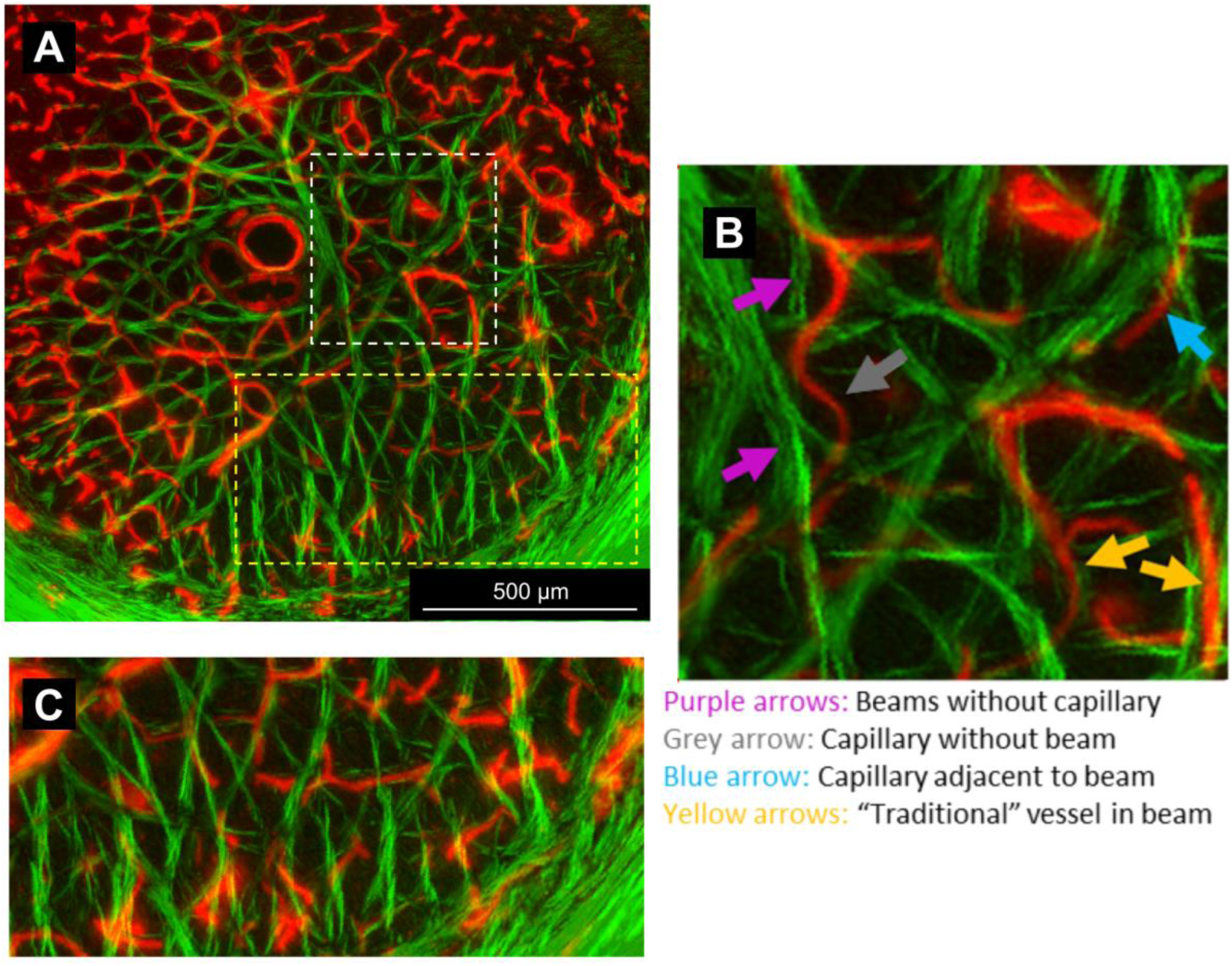
Red (vessels) and green (collagen) merged image of 2 sections (∼32 µm, left). The enlarged region of interest (right) shows 4 different interactions between collagen beams and vessels. Purple arrows: beams without any vessels. Grey arrow: vessel crossing a pore without any beam support. Blue arrow: vessel running adjacent to a beam and even crossing perpendicular to the next beam. Orange arrows: “traditional” vessel within the center of a collagen beam. A) View of the LC region in these sections, B) detail of white box in A, C) detail of yellow box in A. Dye brightness varied. Whilst it may appear as if the yellow box region in panel A has fewer vessels, adjusting the vessel red channel, panel C, it is readily discernible that this is not the case. No significant portions of dye-free LC were observed, indicating complete or near-complete vascular perfusion.

**Figure 7.**
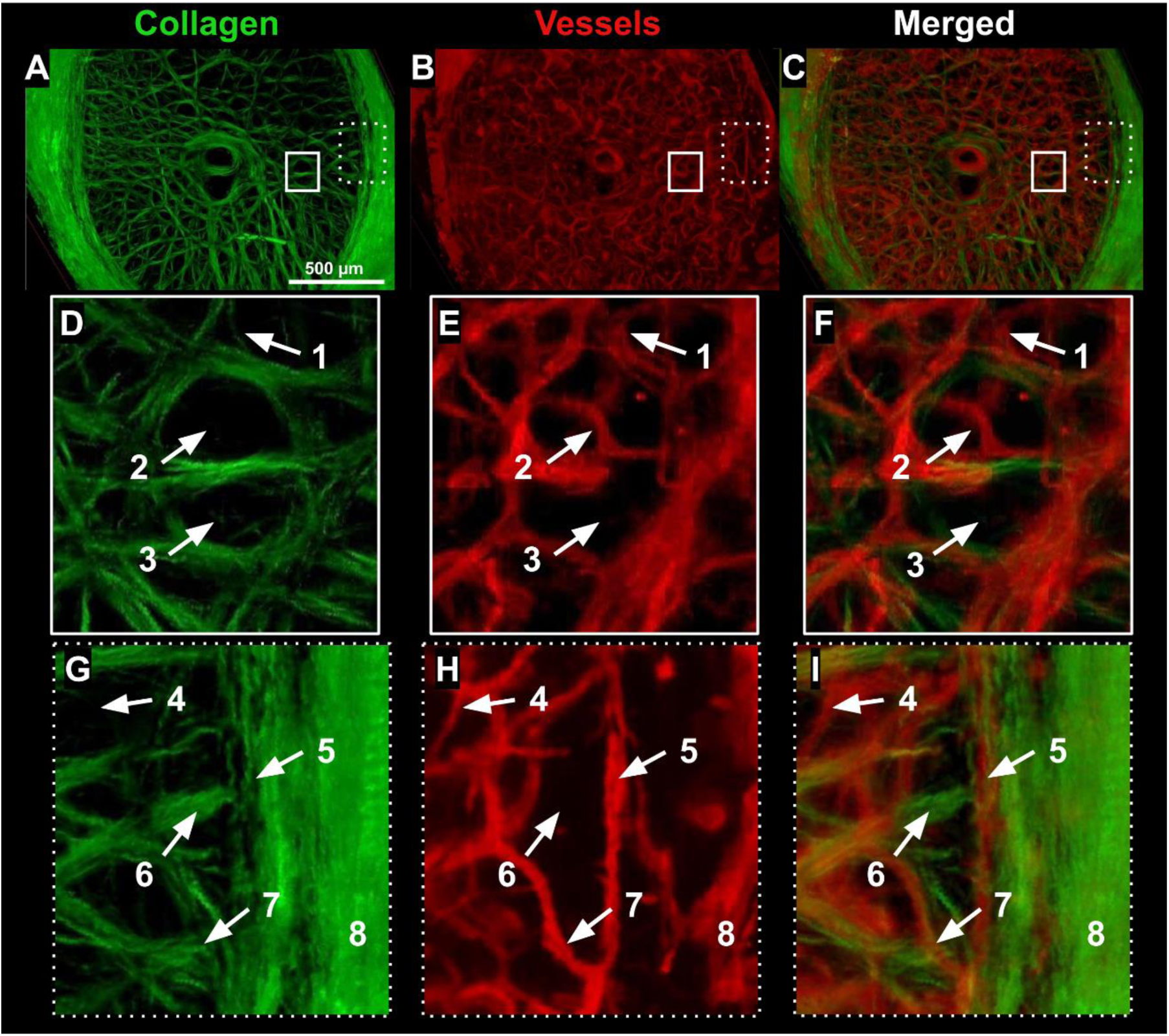
Vessels (DiI in red) and collagen (green) in a monkey ONH. Images are maximum intensity projections of an approximately 256µm-thick region containing the LC. Two areas are highlighted in A-C (solid and dashed boxes) to provide more detail. D-F show close-ups of the central LC within the solid white boxes in A-C. G-H show close-ups of peripheral LC and sclera within the dashed boxes of A-C. Some noteworthy features: 1 & 4) vessels without beams; 2) vessel crossing LC pore; 3) LC pore with no vessels; 5) vessel circling the scleral canal adjacent to the LC; 6) collagen beam without a vessel; 7) vessel feeding into the LC; and 8) vessel circling the canal further into the sclera.

**Figure 8.**
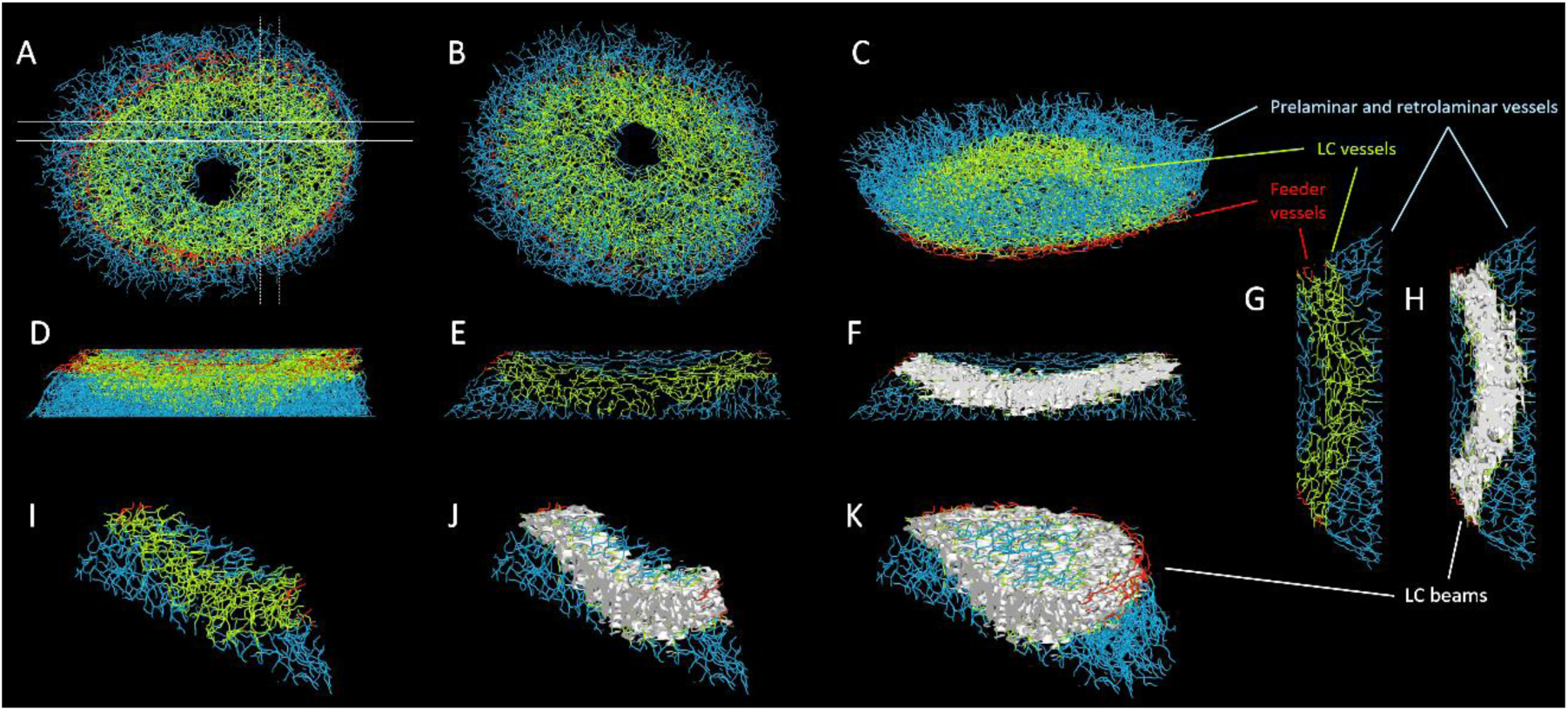
Visualization of the 3D vessel and beam networks. Vessels and beams were segmented in FM and PLM images of every coronal section of a stack that included the LC region. The vessels and beams were reconstructed in 3D over the whole region. The beam segmentations were then used to identify the collagen-rich LC region. Based on the LC region, we then separated the vessels into four groups, shown in this figure using different colors: LC vessels were the vessels within the LC region (green), feeder vessels were peripheral to the LC, connecting to the larger arterioles surrounding the canal (red), and prelaminar and retrolaminar vessels were those within the scleral canal, anterior or posterior to the LC (light blue). The LC vessels were then further subdivided for analysis into those inside or outside LC beams (see **Figures 9, 11, and 12**). The panels in this figure show visualizations of the 3D vessel and beam networks. A-D show views of all the vessels reconstructed. A: coronal view from the front B: coronal view from the back, C: isometric view slightly posterior where the central LC and feeder vessels are clearly discernible. D: longitudinal view from the temporal side. The feeder vessels are discernible but the density of vessels obstructs seeing the LC clearly. To improve the visualization of the LC curved profile, longitudinal slabs were “cut” of both vessels and beams. E-F show a slab cut in the superior-inferior direction from the temporal side. G-H show a slab cut in the nasal-temporal direction seen from the inferior side. I-J show the nasal-temporal slab in isometric view. E, G, and I show only the vessels within the slabs. F, H, and J show the LC beams. The curved shape of the LC is easily discernible in the green LC vessels and white LC beams, surrounded by prelaminar and retrolaminar vessels and a few segments of feeder vessels. K is similar to J, but only one “cut” is shown, leaving the rest of the superior canal region to better visualize the prelaminar and feeder vessels. Note that the slabs are flat cuts of what are complex 3D architectures and therefore some beams and vessels may appear discontinuous when they are, in fact, continuous. To simplify visualization and allow focusing on the characteristics of the vessel network, all vessels are shown with the same diameter, despite the diameters varying. Also, the vessel location and coloring are set according to the location of vessel centroid nodes relative to triangles forming the beam surfaces. As such, they are approximate and some segment ends may extend with a given beyond the region they indicate. The analysis of LC beam and vessel inter-relationship in this work was based on the data as segmented, as described in the manuscript, and is much more precise than the visualization.

Full views of the complete 3D reconstructions of the LC vessels and beams of monkey 2 OD from the front are shown in **Fig. 9**. These images confirm that there were no regions without vessels or beams and illustrate that the LC beam and vessel networks are extremely complex. Visibility through pores depends upon LC orientation, as shown in **Figure 10**. To help visualize the interrelationship between vessels and beams the vessel segments are shown colored according to whether they are inside (blue) or outside (red) a LC collagen beam. Only vessels within the LC region are shown. Compare with the images showing all the reconstructed vessels (**Fig. 8**). The LC beams are shown semi-transparent to allow discerning better the vessels within beams.

**Figure 9.**
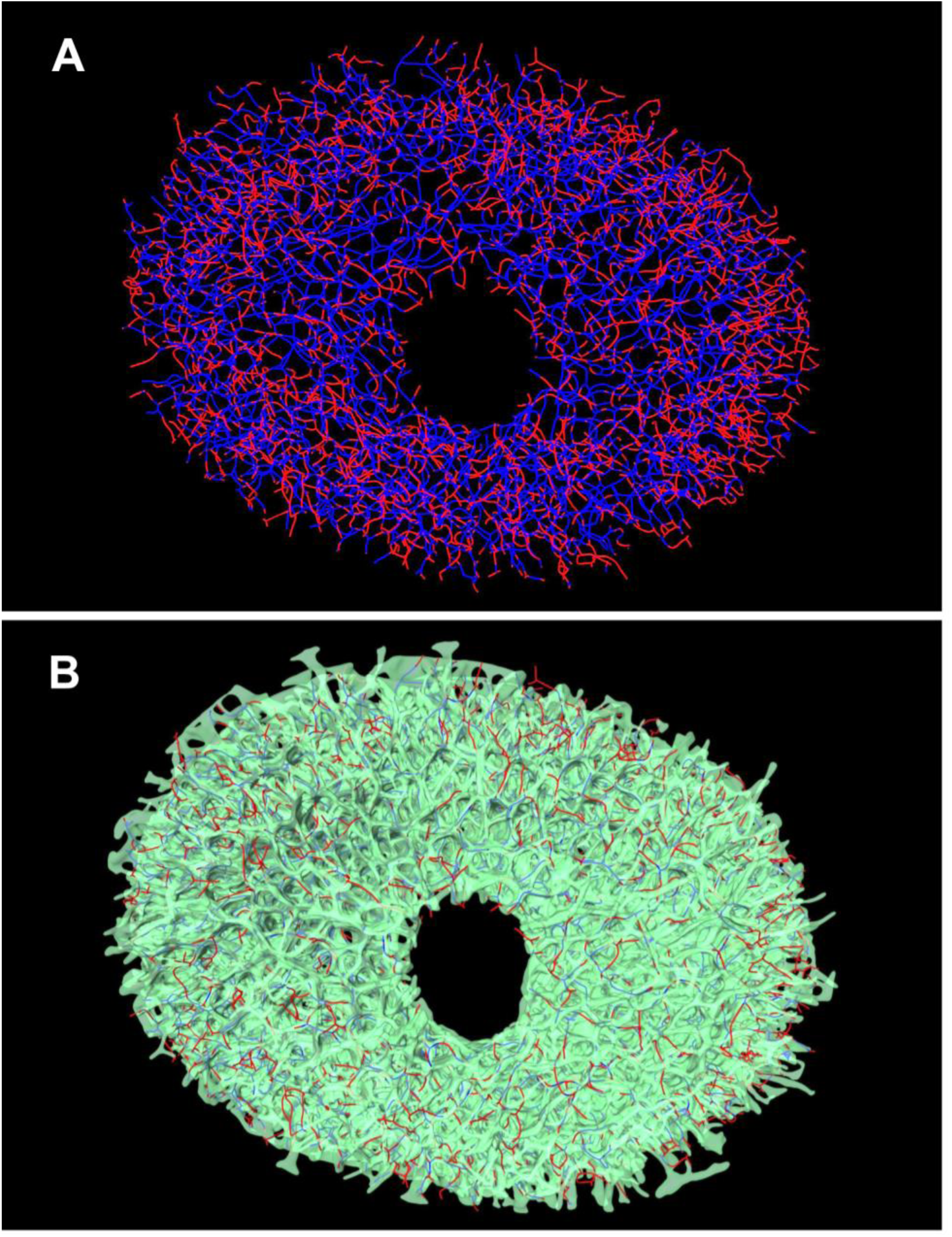
Full views from the front of the 3D reconstructed LC vessels and beams from monkey 2 OD. A) LC vessel segments colored according to their location relative to the beams: blue if inside a beam and red if outside. B) The LC beams are shown semi-transparent, with the LC vessels in red/blue as in Panel A. We were able to reconstruct the collagenous LC beams which vascular corrosion cast techniques destroy during processing. As in **Figure 8**, to simplify focusing on the network, all vessels are shown with the same diameter. Only vessels within the LC region are shown. The extreme complexity of the beam and vessel networks is readily apparent, with no clearly discernible pattern for either or for their inter-relationship. As for **Figure 8**, the segment coloring is approximate, with a few short blue segments that may appear to briefly step outside beams, and red segments within beams.

**Figure 10:**
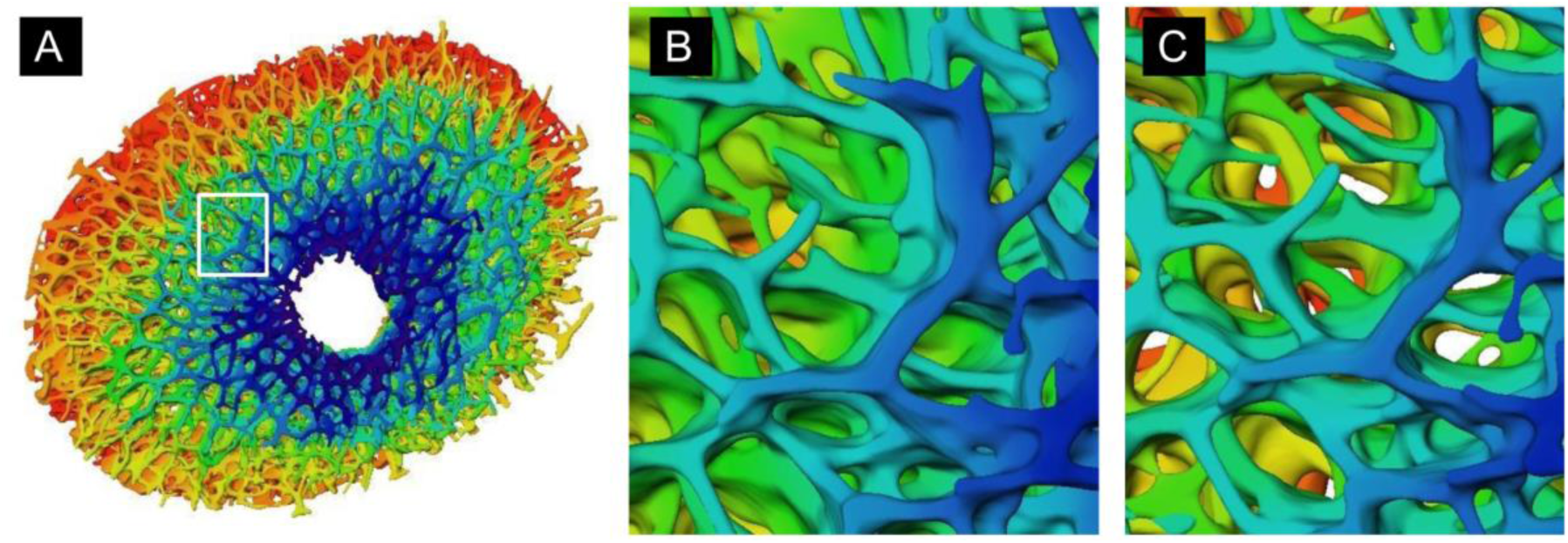
Visibility of pores through the LC depends upon LC orientation. The LC beam surface, color-coded by depth (red is most anterior, blue most posterior). **A** shows a full view of the LC. **B** and **C** are close-ups of the same location differing only on the perspective or direction of view. The colors are useful to discern how deep into the LC one can see. The view in **B** shows no white background, indicating that from this perspective there are no straight paths through the LC. Even a small rotation, as shown in **C**, reveals several points with straight paths through the LC. This illustrates the complexity of the LC and that not seeing all the way through does not mean that there are no pores. Note that because the pores are not straight, it is possible that there are no perspectives in which there is a direct path through the LC. The axon bundles can follow tortuous paths

Magnified views of 3D reconstruction detail are shown in **Figures 11** and **12**. There were an appreciable number of beams with vessels (blue) and beams without vessels (red). Their spatial relationship did not follow any readily apparent pattern. As described in the Methods, we quantified the percentages of *vessels outside beams* and *beams with vessels* in each section of each eye (**Fig. 11** and **Supplementary Table 1**). The number of histological sections analyzed from each eye is included in **Supplementary Table 2**. All LCs sectioned were within a standard range of thicknesses for the monkey eye^44^. *Vessels outside beams* for all eyes was 20.9 ± 12.6%. The mean (± S.D.) percentage of *beams with vessels* was 22.0 ± 7.4%. Neither the percentage of *beams with vessels* nor *vessels outside beams* varied significantly between contralateral eyes of individual monkeys (p = 0.15 and 0.36, respectively). However, greater variability existed between individual monkeys. The percentage of *vessels outside beams* but not *beams with vessels* was significantly different between individual animals (p = <0.0001 and 0.15, respectively). Across all eyes, the percentage of *beams with vessels* was not significantly different from the percentage of *vessels outside beams* (p = 0.46).

**Figure 11.**
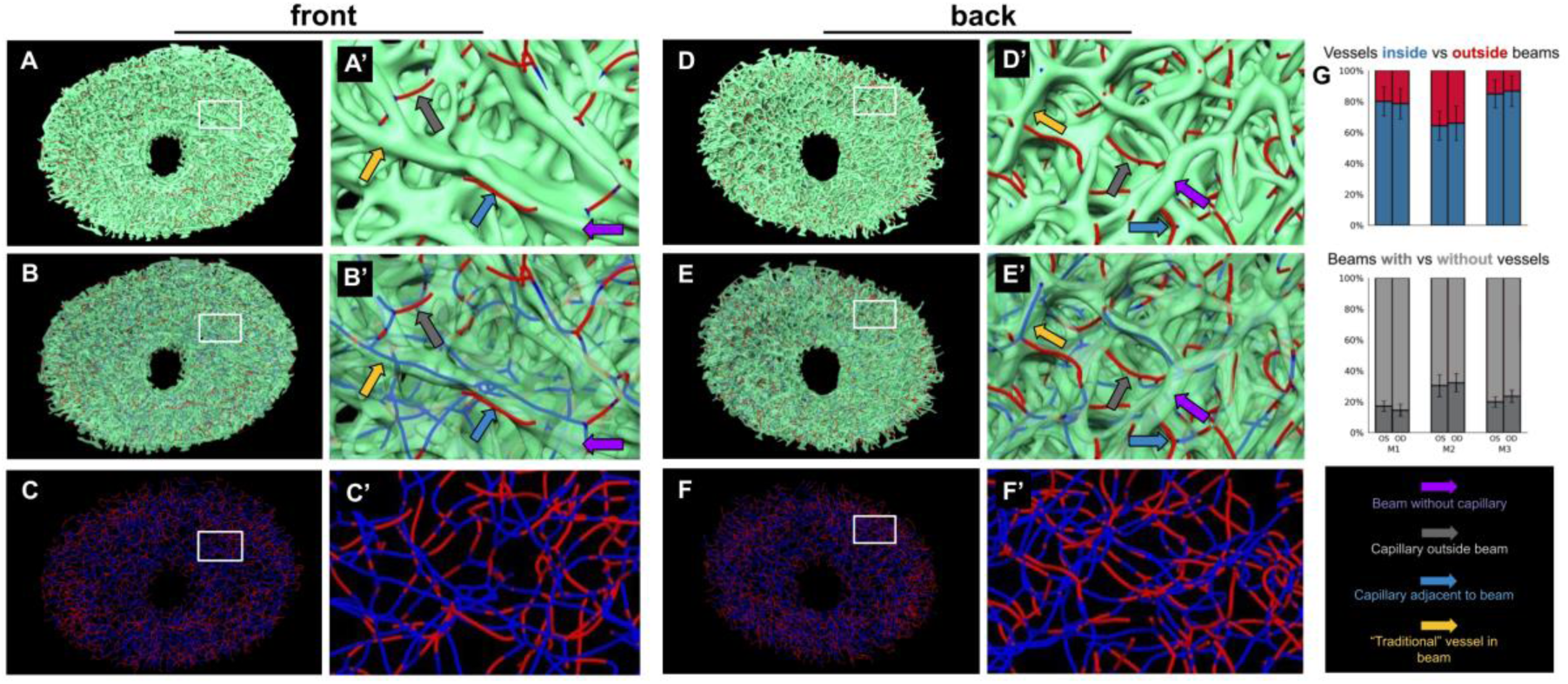
Illustration and quantification of LC vessels inside vs. outside beams. Full views of the full LC of monkey 2, OD are shown from the front in A, B and C, and from the back in D, E and F. A magnification of the rectangle region marked in A, B and C from the front is shown in A’, B’ and C’. Similarly, the rectangle in D, E and F is shown magnified in D’, E’ and F’, from the back of the LC. The first four columns show LC vessels colored according to their location, red when outside beams and blue when inside a beam. Panels in a given column show the same region, varying only on how the beams are shown. Top row (A, A’, D and D’) shows LC beams as solid. This emphasizes LC vessels outside beams (in red). We have indicated a few vessels outside beams (grey arrows) or vessels adjacent to a beam, still outside (blue arrows). As elsewhere, the coloring is approximate and the endpoints of some vessels that are within beams are discernible. The middle row (B, B’, E and E’) show beams semi-transparent. This allows discerning vessels within the beams (blue). We have indicated a few vessels within beams (yellow arrows) and a few beams without vessels (purple arrows). The bottom row (C, C’, F and F’) do not show beams. G. The inter-relationship between LC vessels and beams was quantified, as described in the text. Results for 6 monkey LCs are shown as percentages. The bars in red/blue show the percentage of vessels inside vs. outside beams, whereas the grey bars show the percentage of beams with vs. without vessels. M1: monkey 1, M2: monkey 2, M3: monkey 3. Error bars are the standard deviation over the sections.

**Figure 12.**
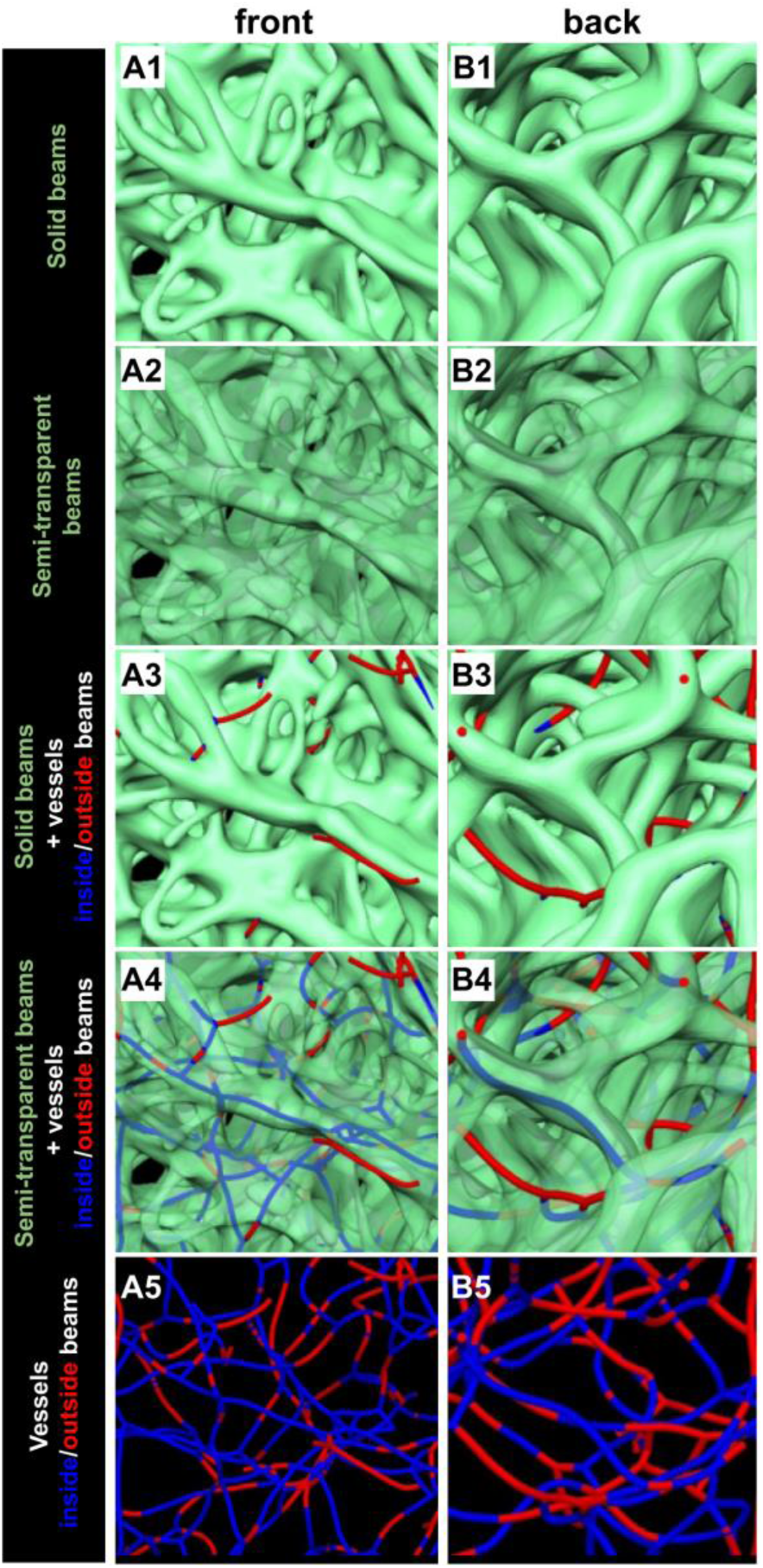
Magnified views of LC 3D reconstructions to illustrate the complex diversity of vessel-beam interrelationship. Two LC locations are shown, one seen from the front (left column, As) and another seen from the back (right column, Bs). Rows show the same location, varying in how the vessels and beams are shown, similar to **Fig. 11**. A1 & B1 show solid LC beams in teal. The back region exhibits slightly wider beams, but overall they are similar. A2 & B2 show the LC beams semi-transparent to enable visualizing the complexity of the structure in depth. A3 & B3 show solid LC beams and LC vessels. LC vessels outside beams are red, and inside beams are blue. A4 & B4 show semi-transparent beams with vessels inside/outside beams shown in blue/red, respectively. Vessels along beams are clearly discernible in both regions. A5 & B5 show only the LC vessel segments, colored as above. No clear pattern of vessels inside/outside beams emerges.

The collagen beam network and the capillary networks are distinct from each other, but there is no evidence of multiple capillary networks. In **Figures 9, 11, and 12**, we show vessels color-coded to indicate whether they are inside or outside beams. Both vessels inside and outside beams show good continuity and interconnectivity with each other as would be expected from a single vascular network. To further help readers grasp the interrelationship between beams and vessels within the LC, we prepared the images shown in **Fig. 13**. The images show the LC collagen beams colored according to local proximity to a vessel inside a beam. Thus, a beam segment with a vessel appears blue and a beam segment that does not have a vessel inside appears read. Note that these plots are different from maps of beam distance to a vessel. We did this because our goal was to illustrate which beams have vessels and which beams do not. If we had used beam distance to a vessel, a vessel adjacent to the beam would have caused the beam to appear blue. Distance to vessels is likely important to perfusion and oxygenation, which is outside the scope of this work.

**Figure 13.**
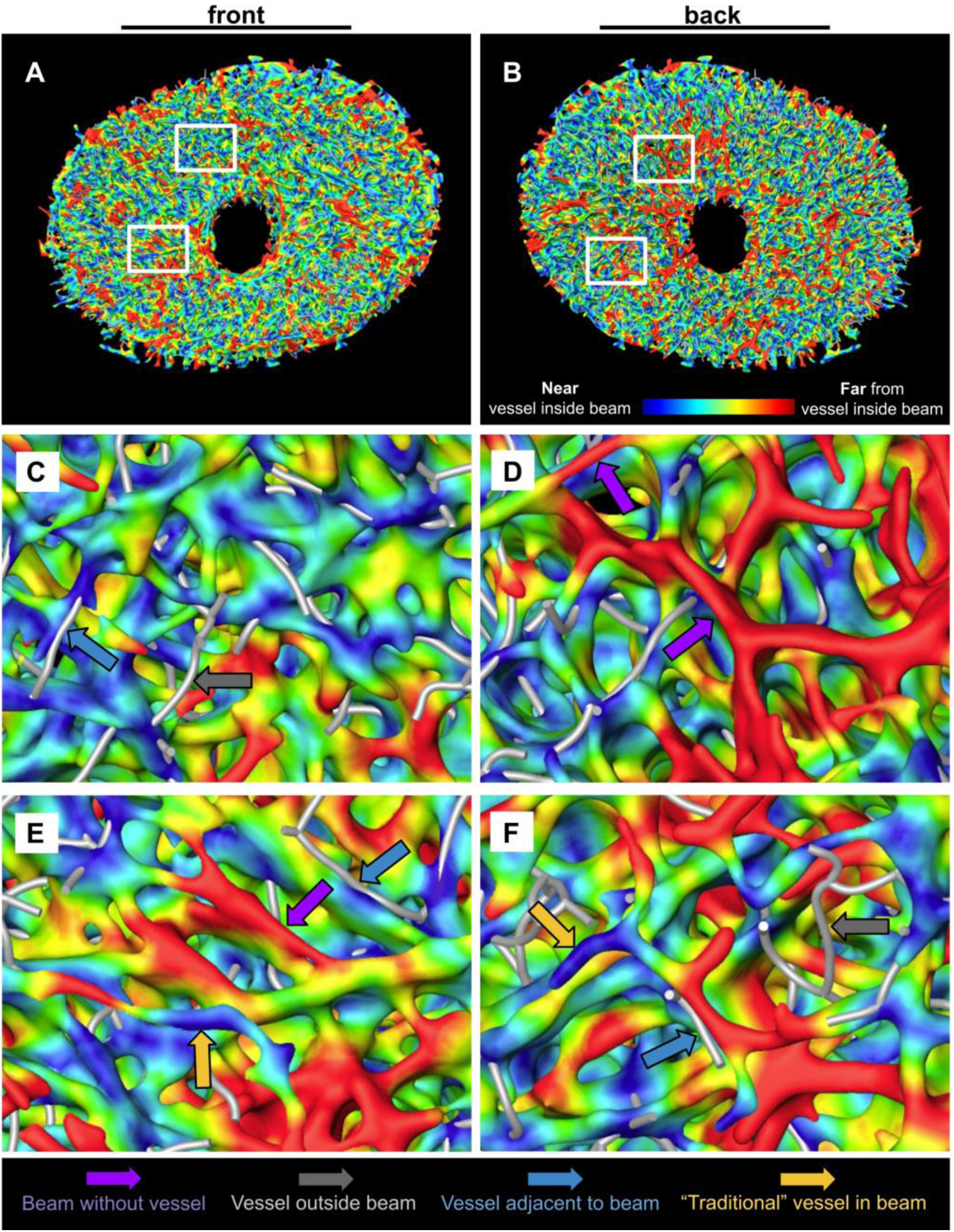
3D reconstruction of LC beams from monkey 2 OD, pseudocolored to indicate proximity to a vessel inside a beam. Reconstructions are shown from the front (A) and from the back (B). Details from the top and bottom boxes indicated in A and B are shown in the second and third rows, respectively. The intent of these images is to help readers grasp the inter-relationship between LC vessels and beams. Where a beam has a vessel within, the beam will be blue. Where a beam does not have a vessel within, it will be red. We have added a few arrows to note examples of beams without vessels (purple arrows), vessels outside beams (gray), vessels adjacent to beams (blue), and “traditional” vessels in beams (orange). These heatmaps are a tool for visualization and do not directly reflect our methods for quantitative analysis.

## 4. Discussion

Our goal was to understand the spatial relationship between the vasculature and the collagen beams of the LC. We quantitatively determined both the global and local relationships of the collagen beams and vessels within the LC of six monkey eyes. Specifically, we quantified the number of collagen beams that contained vessels, collagen beams that did not contain vessels at all, vessels located inside collagen beams, and vessels located outside collagen beams. These data were obtained by examining the ocular tissue after perfusion of DiI and DiD through the carotid arteries of the monkey heads and imaging 16 µm-thick sections with both FM and PLM techniques. From these images, we have made several findings that challenge conventional descriptions of the ONH vasculature. First, we observed that the collagen beams and vessels had distinct topologies. Second, a significant number of LC beams did not contain vessels. Third, a significant number of LC vessels were not contained within beams. These findings were consistent between contralateral eyes with larger variations found between individuals. We are not aware of other quantitative reports of the interrelationship between LC vessels and beams. The implications of these findings are discussed below.

### LC collagen beams and vessels have distinct topologies

We noticed that the vessels weaved in and out of the collagen beams and were not always located within the center of the collagen beams. This demonstrates that the common perception of every vessel being contained within a lamina beam is not entirely accurate. The occurrence of disparate networks, marked by the observed *beams without vessels* and *vessels outside beams*, is likely due to the different roles that they each play in supporting the axons that pass through the LC. The collagen beams function to provide mechanical support to the axons whereas the vessels in this region function to provide oxygen and other nutrients. The optimal structure for one may not be the optimal structure for the other. Further exploration of these topological relationships in both healthy and pathologic eyes may inform novel risk factors for ONH disease.

### Many collagen beams do not contain vessels

This is likely due to the vast differences in the topologies of the two networks. Because there was a significant number of *beams without vessels* within them or along their perimeter, we speculate that these beams play a larger role in mechanical support for the LC. The presence of collagen beams that do not contain vessels suggests that the collagen beam network is designed to provide the maximum mechanical support to the axons as they pass through this region, independently of vascular demand. Interestingly, these numbers are also conserved between the contralateral eyes.

### Some vessels do not reside within a beam

We observed *vessels without beams* that ran alongside or adjacent to a collagen beam, crossed perpendicular through a beam, and even spanned across the pores between collagen beams. This variation likely serves the benefit of allowing axons easier access to oxygen and other necessary nutrients. These vessels are likely encased with astrocyte end feet, as is other microvasculature in the central nervous system^45^. However, we have not analyzed this specifically. Additionally, *vessels without beams* may be able to make more contact with resident astrocytes and pericytes, allowing for effective neurovascular coupling. Although the consequences of having *vessels without beams* remain unclear, future studies should investigate whether these vessels are more at risk to changes in IOP than the vessels located within a collagen beam.

Both PLM and DiI perfusion/FM are well-established methods for detection of collagen and vasculature, respectively. Nevertheless, it is useful to consider how potential false negative detection of collagen and vessels may impact our results. If we were unable to detect collagen around some vessels, we could erroneously think that a vessel is not in a beam. We believe that this is extremely unlikely as it goes against the very substantial body of literature demonstrating evidence of collagen birefringence. Specifically, there is strong support for the ability of PLM to visualize all collagen present within tissue sections, including the lamina cribrosa^27,32,46,47^. No differences in collagen abundance or distribution were observed between results from PLM and results from OCT^20,48^, SHG^49,50^, or in samples stained to mark collagen^51^. Further, our 3D reconstructions show complete, well-formed beams, similar to those observed with other techniques^52^. Our 3D reconstructions of LC collagen show similar pore architectures to those created by others from human^53^ and monkey eyes^34^. Our analysis of sensitivity showed that variations in beam and/or vessel width have only a slight effect on the quantitative results, and thus do not affect the main conclusions in the paper. Comparison of beam widths between PLM and other techniques revealed excellent agreement, without systematic differences, and variations that are substantially smaller than the range considered in the sensitivity study. Our choice of technique, namely PLM, did not directly determine our findings compared with those had we imaged all samples using picrosirius red or SHG. However, these results suggest that equivalent conclusions would be found if the work had been carried out using picrosirius red or SHG imaging for LC beams.

For all the LC vessels to be enclosed in beams, the LC would have to be extremely dense, with beams and pores that look totally different. Regarding false negative detection of vessels, it is possible that some vessels within were not visualized, for instance in the case of incomplete DiI perfusion. Although we cannot claim that our method is guaranteed to have perfused every single vessel, the relative uniformity of fluorescent signal distribution in the LC provides evidence that much of the vasculature has been perfused with DiI/DiD label. By following the labeled vessels we are able to identify a vast continuous vascular network. We do not observe any regions with clearly missing vessels, or unexplained end-points. Most importantly, the potential for missing some beams and/or some vessels within the LC would not change the three main conclusions of the manuscript stated above.

Conversely, we also consider the potential of false positive detection of collagen and vessels. For similar reasons stated as above, although it is expected that some tissues within the ONH exhibit some birefringence (such as axon microtubules^54^), widespread false positive detection of collagen from PLM is very unlikely. Nevertheless, if we had considered some other birefringent constitutive component of the tissue as collagen, this would likely appear as part of the collagenous LC beams. As for false positive detection of vessels, this is one of the main reasons for us to use intravascular dye perfusion. This technique is very well established as reliable^27,32,46,47^. This is also the method of choice for other state-of-the-art studies of ONH vasculature^55^. Detecting false vessel positives would require that something else is fluorescent in the same wavelengths, despite having no labels. This was easily disproven by imaging negative controls (results not shown) which did not demonstrate any appreciable vascular-specific signal. Another possibility is that the dye may have leaked through a burst vessel. From our pilot studies, we observed that these leaks look quite different from vessels and are easy to distinguish when they are larger than a few pixels. Hence, we are confident that our segmentations do not include them. In any case, the false positive vessels would have to be substantial in quantity to affect our main conclusions that LC beams and vessels have distinct topologies. A few “fake” vessels would not be enough.

Within recent decades, many studies of ONH vasculature have been conducted in vivo utilizing non-invasive imaging techniques such as fluorescein angiography^56,57^ and optical coherence tomography angiography.^58^ Each of these techniques, while powerful tools for answering many questions about superficial vasculature, suffer from limited depth penetration and shadow artifacts. These limitations preclude high-resolution imaging of deeper vascular plexuses such as those in the LC and retrolaminar tissue.^19,59,60^ Other techniques, including optoacoustic, ultrasound, MRI, and micro-CT have excellent signal penetration but lack the spatial resolution required for detailed analysis of the LC.

We utilized a combination of perfusion labeling, serial histology, and 3D reconstruction. In some ways, this combination of methods is a return to the more traditional approaches used during the 1950s to 1980s. However, we have the substantial advantage of more advanced microscopy and computational resources to reconstruct high-fidelity 3D vessel maps across large regions that could not have been obtained previously. Our approach also has several other advantages over commonly used methods for studying the ONH vasculature. For instance, one of the more commonly used methods has been transmission electron microscopy (TEM).^61,62^ TEM can provide excellent high-resolution images, but the field of view is too limited.^63^ Scanning electron microscopy provides excellent resolution and field of view, but usually provides little information about the sample interior.^64^ Creating vascular molds of the ONH requires supraphysiologic pressures for perfusion of viscous casting agents.^65–67^ As vascular casts require digestion of the tissue after these perfusions, this method is not well-suited for understanding the relationships between vasculature and its surrounding extracellular matrix. Additionally, tissue clearing has served as an excellent way to visualize vasculature within intact tissues such as the brain.^68,69^ Casting methods often require the use of solutions that can shrink, expand, or otherwise distort tissues which may affect some interpretations. In contrast, our approach allows for low-viscosity aqueous solutions to be perfused through the central retinal artery and ciliary arteries at controlled, lower pressures. With our methods, vasculature was visualized at high resolution while extracellular matrix was preserved. These advantages allowed us to generate 3D geometries throughout the complete LC, uniquely well-suited for analysis of intricate vessel-beam interactions. For these reasons, the approach developed for this study is advantageous.

In the study detailed in this paper, we aimed to identify all potential vessels. To accomplish this goal, we perfused eyes with vascular labels at different IOPs, spanning a wide, but still physiologic, range of 5 to 25mmHg, as detailed in the **Supplementary Figure 1**. We then created composite images via union of all labelled vasculature, regardless of the IOP at which they were labelled. We selected DiI and DiD to label vasculature due to their extensive documentation in the literature as reliable vascular markers.^23–25^ Marking vascular endothelium with labels such as CD34, vWF, and lectins, can require multiple steps. Signal intensities achieved by these methods are lower and often more variable than that achieved with DiI/DiD perfusion labelling.^23^ It is possible, although to the best of our knowledge unproven, that some of the LC vasculature is only “active” within a specific IOP range. Combining the vessels prevented us from discerning potential effects of IOP, which is of great interest to understand eye pathophysiology, and in particular glaucoma, and should therefore be the focus of future research. Nevertheless, despite differences in IOPs between contralateral eyes, we observed low contralateral variability in the relative abundance of beams with/without vessels and vessels inside/outside beams. This may indicate that the level of IOP did not affect the interrelationship between beams and vessels, and thus that these choices did not affect our main conclusion that LC vessels and beams have distinct topologies.

This manuscript is not intended as a comprehensive review of the literature on structure and ultrastructure of the monkey or human ONH. We direct readers interested in more information on this topic to a number of excellent sources.^5,14,61,70–72^ Unlike others before it, this manuscript places the focus on the interrelation between vessels and collagen beams. Additionally, many of these studies were conducted decades ago, and technological advances now allow for high-resolution digital 3D reconstructions that are several gigabytes in size which were simply impossible before. This facilitates visualization and quantification of information that could be misinterpreted when viewing single sections or projections.^40^

Our methods have potential limitations. One issue relates to any type of *ex vivo* vessel perfusion study because the delay between death and perfusion leads to intravascular clotting that can block perfusion. We were able to obtain monkey heads within minutes of sacrifice and begin the perfusion process via the carotid arteries within an hour of sacrifice, which likely mitigated this concern. Furthermore, we used ample PBS flushing to remove blood from inside these vessels. Even if a partial clot remained, the vessel downstream could still be labeled. Hence, while a blockage is still possible, we believe that the vast majority of vessels were labeled and that no large areas were affected. This is also supported by the close similarity in numbers between contralateral eyes. Finally, although clotting could change the exact numbers measured, several of the most important findings would not be affected. For instance, that there are a substantial number of vessels outside collagen beams.

We analyzed tissues that had been processed histologically. Thus, there may have been artifacts as a result of the fixation or sectioning, including tissue distortion or shrinkage. However, we have shown previously that our method of formalin fixation has minimal effects on size and shape of ocular tissue.^26,46^ Future work could use fiducial markers to correct for any tissue warping during sectioning.^73^ Because each eye was treated in the same way, comparing the vessels and collagen beams from all eyes to one another is appropriate. In addition, our cryosection samples for structural studies were 16 µm-thick which resulted in a more limited depth resolution compared to the in-plane resolution. While serial sectioning has been proven to be an effective approach in many studies, it may introduce some quantification errors due to its relatively low depth resolution. Both collagen beams and vessels are organized in 3D and a section of 16 µm may separate a single beam or vessel into two or more sections. Thus, the ratio measurements may be either under- or over-estimated locally. Across the entire section, however, the overall effects due to sectioning are to some extent averaged out and thus the potential impact for the entire section is less severe than any local effect.

In future studies, a higher depth resolution is desired which will allow for higher fidelity 3D reconstruction of the collagen beam and vessel networks, potentially leading to 3D quantifications. Techniques like a tape transfer system can significantly reduce the minimum section thickness to as low as 2 µm and could be implemented in future studies. In this initial study, we used a binary classification to define vessels as being within or not within a beam. Our algorithm was intended to help test the one-to-one model of beams and vessels. Similarly, we defined beams as having or not having vessels and vessels as either inside or outside a beam. From the images collected in this study, it became apparent that vessels can cross beams, be parallel to beams, or wind in and out of beams. It will be important for future studies to employ a more refined characterization of the relationships between LC beams and vessels to further describe and quantify the newly found complexities of these relationships that we identified. Such an algorithm should be able to distinguish, for example, if a vessel is within a beam by traversing it longitudinally, or if it crosses at a relatively large angle, which the algorithm used in this current work could not do.

In conclusion, to the best of our knowledge, we have presented the first 3D data of the vessels and collagen beams within the monkey LC. Surprisingly, we discovered that the collagen beams and vessels had vastly different topologies and networks across the LC. In addition, we provide evidence that a significant number of collagen beams do not have a vessel inside of them, and that a significant number of vessels are not located within a collagen beam. This variation is likely due both to the demand for carefully tailored structural support as well as the high metabolic demand of the axons within the LC. These findings emphasize the critical need to better understand the role of LC vessels in the normal physiological state, in aging, and in important optic nerve diseases. Specifically, and with respect to the latter, with information derived from this study, we now have a greater understanding of the potential vascular risk factors associated with glaucoma and NAION, two important and relatively common forms of blindness.

## Disclosures

B.L. Brazile was at the University of Pittsburgh when he contributed to this work. He is now at Baxter; B. Yang, None; S. Waxman, None; P. Lam, None; Dr. PY Lee, None; Y Hua, None; Dr. Voorhees was at the University of Pittsburgh when he contributed to this work. He is now at Johnson & Johnson; A.L. Gogola, None; J.F. Rizzo, None; T.C. Jakobs, None; I.A. Sigal, None

## Acknowledgments

We gratefully acknowledge support from National Institutes of Health R01-EY031708, R01-EY023966, R01-EY028662, P30-EY008098, T32-EY017271, Eye and Ear Foundation (Pittsburgh, Pennsylvania), Research to Prevent Blindness. Matthew Smith and Samantha E. Schmitt for help with the perfusion techniques. For providing the tissues Steven D. Abramowitch, Pamela A. Moalli and Stacy Palcsey, supported by NIH R01-HD045590.

## Notes

### Summary of Updates

Add two authors that were inadvertently missed in the previous submission (Hua and lee).

