## Supplementary for "Lamina cribrosa vessel and collagen beam networks are distinct"

**Short Title:** Lamina cribrosa microvasculature

* **Correspondence**:
Ian A. Sigal, Ph.D.
Laboratory of Ocular Biomechanics
Department of Ophthalmology, University of Pittsburgh Medical Center,
203 Lothrop St. Rm. 930, Pittsburgh, PA, USA. 15213
 www.OcularBiomechanics.org

**Keywords:** Collagen, Vasculature, Lamina Cribrosa, Optic Nerve Head, Morphology

**Disclosures:** B.L. Brazile was at the University of Pittsburgh when he contributed to this work. He is now at Baxter; B. Yang, None; S. Waxman, None; P. Lam, None; Dr. PY Lee, None; Y Hua, None; Dr. Voorhees was at the University of Pittsburgh when he contributed to this work. He is now at Johnson & Johnson; A.L. Gogola, None; J.F. Rizzo, None; T.C. Jakobs, None; I.A. Sigal, None.

**Funding:** National Institutes of Health R01-EY031708, R01-EY023966, R01-EY028662, P30-EY008098, R01-HD045590, R01-HD083383, T32-EY017271, Eye and Ear Foundation (Pittsburgh, Pennsylvania), Research to Prevent Blindness.

| Average percent per section ± SD | Monkey 1 | | Monkey 2 | | Monkey 3 | |
|---|---|---|---|---|---|---|
| | OS | OD | OS | OD | OS | OD |
| Beams with vessels | 17.0 ± 3.4 | 14.4 ± 3.8 | 30.3 ± 7.1 | 32.1 ± 5.9 | 19.8 ± 3.4 | 23.5 ± 4.1 |
| Beams without vessels | 83.0 ± 3.4 | 85.6 ± 3.8 | 69.7 ± 7.1 | 67.9 ± 5.9 | 80.2 ± 3.4 | 76.5 ± 4.1 |
| Vessels outside beams | 21.5 ± 9.9 | 20.0 ± 9.4 | 35.7 ± 9.4 | 34.1 ± 11.1 | 15.1 ± 9.0 | 13.3 ± 10.0 |
| Vessels inside beams | 78.5 ± 9.9 | 80.0 ± 9.4 | 64.3 ± 9.4 | 65.9 ± 11.1 | 84.9 ± 9.0 | 86.7 ± 10.0 |

**Supplementary Table 1.** Summary of the beams with vessels within them and vessels residing outside of beams for each eye.

| Sections analyzed | OS | OD |
|---|---|---|
| Monkey 1 | 16 | 24 |
| Monkey 2 | 15 | 15 |
| Monkey 3 | 29 | 31 |

**Supplementary Table 2.** The number of histological sections in which vessels and beams were quantified from each eye. Note that while sections were each 16 µm-thick, the cupped shape of the LC as well as small deviations in the plane of sectioning from the coronal plane can increase the number of sections per eye that contained LC. For example, 31 16 µm-thick sections containing LC did not equate to a 496 µm-thick LC.

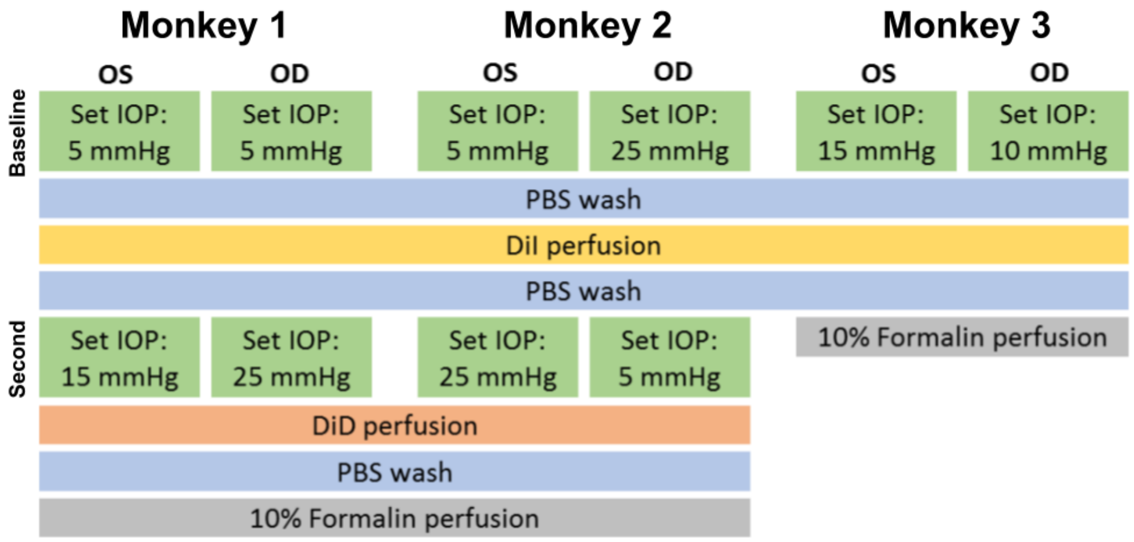


**Supplementary Figure 1**. Schematic diagram of the IOP setpoints and perfusion procedure for each monkey eye used in this study. Monkeys 1, 2 and 3 were 12, 16 and 12 years of age, respectively.

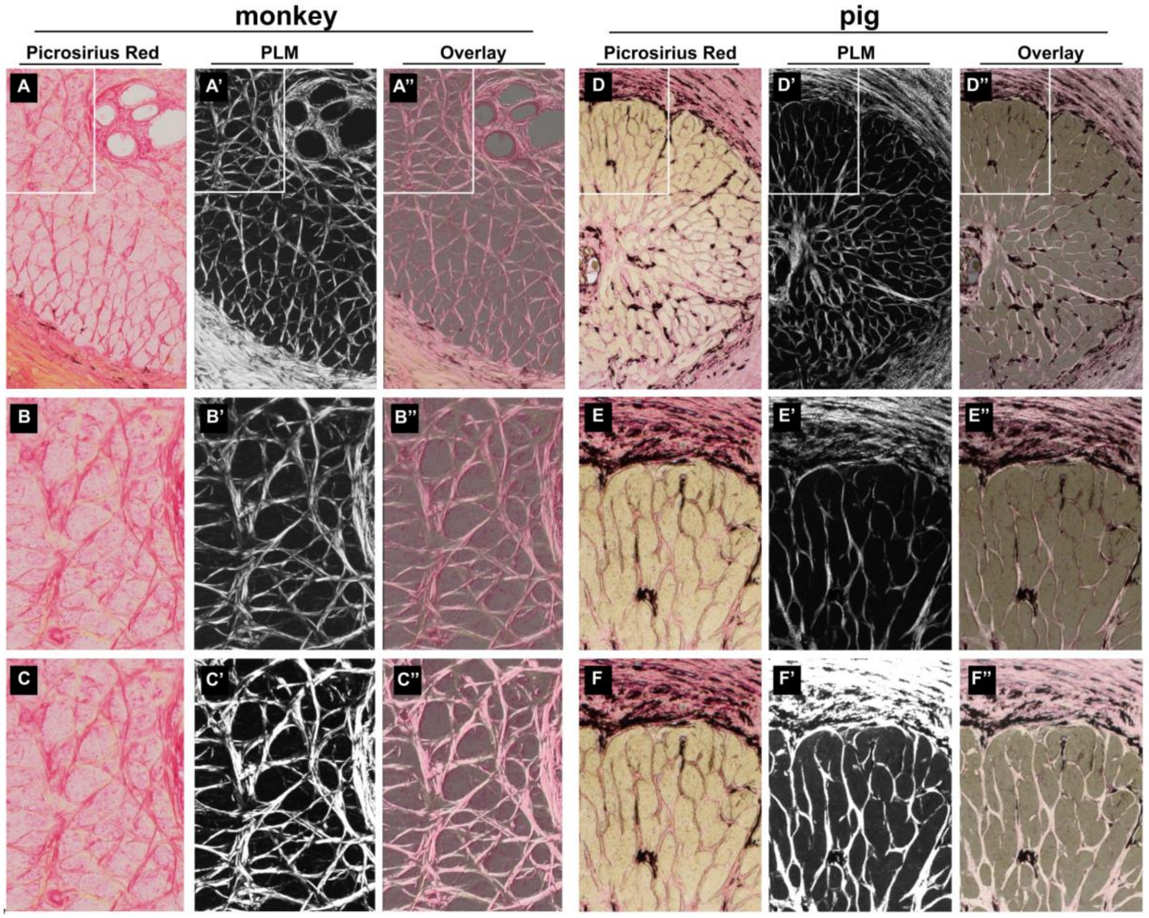


**Supplementary Figure 2**: Polarized light microscopy (PLM) detection of LC collagen beams is in line with other well-established methods of collagen visualization and is consistent across species. A) A 16µm-thick coronal cryosection through the monkey LC was labeled with picrosirius red (PS-red), a collagen stain, and imaged via brightfield microscopy. Collagen is visible as darker pink/red. A') The same section imaged via polarized light microscopy reveals collagen beams, shown in white. These two methods show the same pattern of collagen visualization. This is evident in an overlay of PLM and PS-red images in A''. B-B'' show detail of the region in A-A'' within the white box. It should be noted that exposure of PLM images was optimized to capture as much image information as possible. Thin collagen beams may appear dim or absent under certain viewing conditions. However, these images do contain the information necessary to visualize and reconstruct all collagen present. To best demonstrate that our PLM imaging captures all collagen visualized in the PS-red images and does not show any additional signal in the dark pores, we include panels C-C'' in which PLM image brightness was increased. D-F'' show images captured and organized the same way as for the monkey LC, but for the pig LC. These techniques detect collagen and do not discriminate between the species of the tissue.

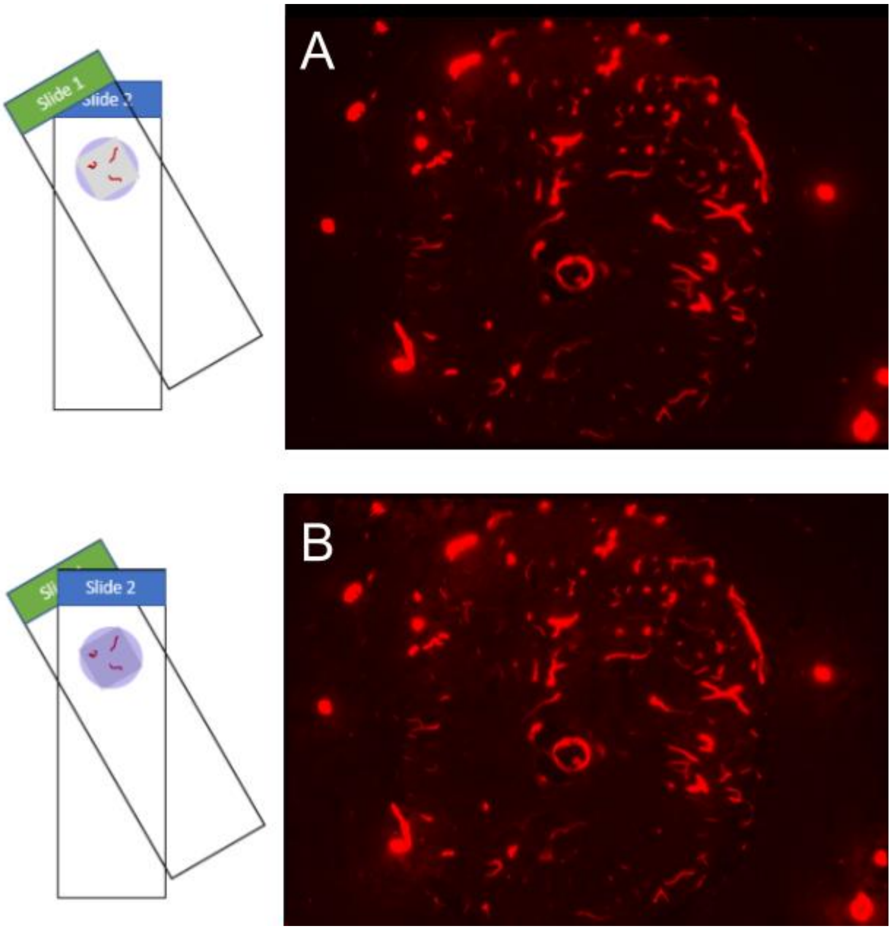


**Supplementary Figure 3.** Evidence that collagen does not obscure DiI/DiD signal.  A 16 µm section of monkey ONH labelled intravascularly via fluorescent DiI/DiD (Slide 1, green) was imaged after being placed on top of a 16µm section of scleral collagen (Slide 2, blue).  Slide 1 was then imaged after the collagen in Slide 2 was placed over top of the sample on Slide 1 (B).  The resulting images demonstrated virtually no differences in apparent vascular labelling.  This demonstrates that presence of collagen as thick as the ONH sections themselves does not impact detection of vessels.
